# DNA methylation clocks struggle to distinguish inflammaging from healthy aging, but feature rectification improves coherence and enhances detection of inflammaging

**DOI:** 10.1101/2024.10.09.617512

**Authors:** Colin M. Skinner, Michael J. Conboy, Irina M. Conboy

## Abstract

Biological age estimation from DNA methylation and determination of relevant biomarkers is an active research problem which has predominantly been tackled with black-box penalized regression. Machine learning is used to select a small subset of features from hundreds of thousands CpG probes and to increase generalizability typically lacking with ordinary least-squares regression. Here, we show that such feature selection lacks biological interpretability and relevance in the clocks of the first- and next-generations, and clarify the logic by which these clocks systematically exclude biomarkers of aging and disease. Moreover, in contrast to the assumption that regularized linear regression is needed to prevent overfitting, we demonstrate that hypothesis-driven selection of biologically relevant features in conjunction with ordinary least squares regression yields accurate, well-calibrated, generalizable clocks with high interpretability. We further demonstrate that the interplay of disease-related shifts of predictor values and their corresponding weights, which we term feature shifts, contributes to the lack of resolution between health and disease in conventional linear models. Lastly, we introduce a method of feature rectification, which aligns these shifts to improve the distinction of age predictions for healthy people vs. patients with various diseases.

**Key Findings:**

- There is no apparent biological significance of the CpGs selected by first- and next-generation clocks
- The range of residuals for first- and next-generation clock predications on healthy samples is very large; for all models tested, a prediction error of +/-10-20 years is within the 95% range of variation for healthy controls and does not signify age acceleration
- There is no significant shift in the mean of residuals for patient populations relative to healthy populations for most studied first- and next-generation clocks. For those with significance, the effect size is very small.
- Hypothesis-driven feature pre-selection, coupled with modified forward step-wise selection yields age predictors on par with first and next-generation clocks. EN/ML is not needed.
- Disease-related shifts at different CpG probes, along with learned model weights, can be either positive or negative; their combination leads to de-coherence effect in linear models.
- Model coherence can be induced by rectifying features to have only positive shifts in patient samples; this provides a better resolution between health and disease in DNAm age models, and expectedly, introduces more non-linearity to the input data.

## Main

It was demonstrated in 2013 by Horvath that probed DNA methylation levels at a subset of CpG sites in multiple tissues can be used as predictors of chronological age [1]. This was done by training a predictive model using elastic net (EN) regression, a penalized least squares regression which performs feature selection [2]. The same year Hannum et al. published a similar model trained on DNA methylation samples in whole blood [3]. These chronological age predictors were dubbed DNA methylation (DNAm) clocks and have become colloquially known as the first-generation clocks.

With little overlap between the CpG subsets selected by the Horvath multitissue and Hannum blood clocks, and with the response variable in both clocks being known chronological age, support for the biological relevance of the clock predictions was lacking. The next generation of DNAm clocks were aimed at identifying subsets of CpGs which can predict biological age or measure biological age acceleration. While the precise methods and training datasets for each next generation clock differ, they follow a general, two-stage construction: an initial filtering, or feature *pre-selection* step whereby CpG probes which are correlates of some physiological or clinical metric of biological age are kept and the rest are discarded, and then using EN to both perform further feature selection from that smaller subset of CpG probes, and to predict either age, time-to-death or some other age surrogate from those selected features [4–7]. It has been shown that DNA methylation clocks—mathematical models with parameters optimized via machine learning—can yield generally accurate age predictions for humans and other mammals, usually by taking linear combinations of individual CpG probe measurements. When regressing model predictions on sample ages, the error, or residual, between the line of best fit and a subject’s actual chronological age has been proposed to indicate an acceleration of biological aging [1, 3–5, 7–12].

However, it remains unclear whether and to what degree such model variance is from random experimental variation and/or healthy epigenetic dynamics versus actual changes in health or biological aging. Epigenetic regulation of cell-fates (myofiber vs. hepatocyte, vs. leukocyte, etc.) needs to be stable, in contrast, epigenetic dynamics is required for healthy cellular and tissue responses to their environments, changing with our daily routine, at night, and even with weather and financial status [13–22]. In out-of-sample data, the differences in healthy dynamics, as well as random and systematic errors, known as batch effects, and different data preprocessing, normalization and other background corrections can all potentially lead to high residuals which would obviously not be attributable to age acceleration. Thus, a reliable metric of epigenetic shifts must be robustly above batch effect noise and normal, healthy dynamics to be broadly useful to clinicians and researchers.

Here, we expand the notion of the low accuracy and poor resolution between health and disease in linear DNAm clocks [23], with novel findings that explain why ML-trained EN predictors underperform in these tasks. Furthermore, our results show that hypothesis-driven feature selection and interpretable ordinary least squares regression can effectively replace black-box machine learning methods and regularization for feature selection. Lastly, we discovered that counterproductive combinations of model weights and disease-related shifts in beta values create a net cancellation effect in a linear model. This causes what we term an *incoherence* effect, where the model fails to distinguish between healthy and patient populations despite actual differential methylation profiles between the two groups. To address this, we developed a method called *feature rectification*, which produces a *coherent* shift effect between features, improving the capacity of linear models to capture methylome differences of aging in healthy vs. patient populations.

## Results

### Analysis of elastic net CpG selection

The elastic net performs feature selection via the L1 norm and minimizing the cost function over the training data [2]. However, in biology, the focus is not on minimizing a mathematical function but on the specific CpG sites and their relationship to changes in gene expression accompanying aging. Therefore, we asked three aging-relevant questions: 1) whether a CpG probe’s correlation with chronological age was related to its importance in the model,

1. if EN selects CpGs based on an increased, age-related dysregulation, and
2. the possibility that the elastic net selects CpGs that are associated with inflammaging.

These studies were applied to all major current clocks: first generation Horvath multi-tissue and Hannum blood age predictors, and the next generation predictors, PhenoAge, AdaptAge, DamAge and CausAge. Additionally, we analyzed the GpG probe selection for DunedinPACE, which is a putative measurement of the rate of biological aging. For our analysis we created a composite dataset (GSE42861, GSE125105, GSE72774, GSE106648) with datasets where DNA methylation was probed from either whole blood or PBMCs from healthy controls and patients suffering from diseases that are associated with inflammaging, such as rheumatoid arthritis, Parkinson’s disease, depression, and multiple sclerosis [24–30].

First, we tested for correlation between a CpG’s importance in a model and the change in its beta values with age. Normalized feature importances were calculated for each CpG of each DNAm clock (methods), with importances near zero indicating low importance, and a value of one indicating the most influential CpG in the model. For each model we regressed the beta values of each CpG individually on the chronological ages of the healthy controls and plotted its R-squared value against its relative importance (**Figure 1a**). Interestingly, in contradiction to the notion of predicting aging from the changes in the methylome, the most important CpGs were, in all models, those unchanged or changed the least in their methylation with age. Given that there are probes for which |*r*| *>* 0.8 for their age correlation (e.g. cg16867657), these results suggest that correlation of CpG methylation with age is not the primary criterion for its importance in a model, otherwise such CpGs would tend to be over-represented in EN models, and with higher importances than CpGs with lower age correlations.

**Fig 1:**
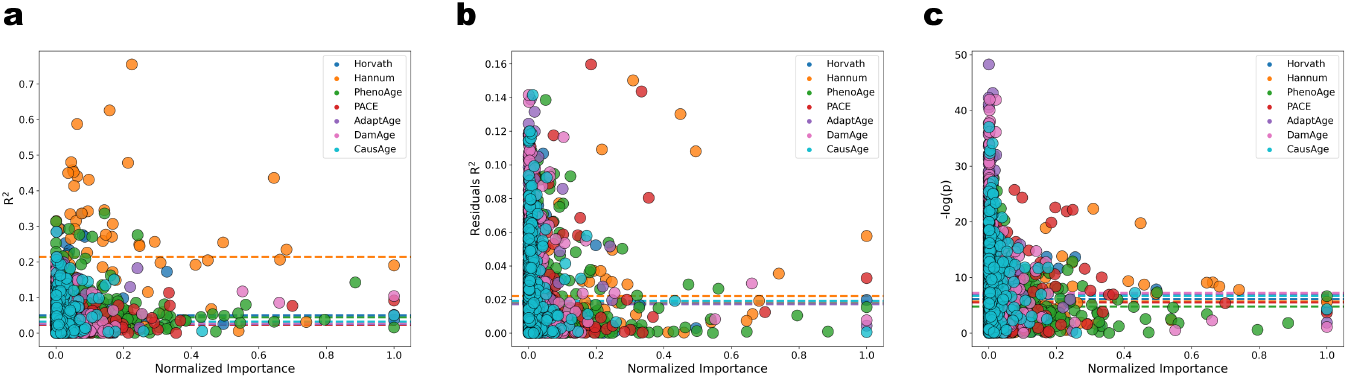
**a** R-squared of the age correlation of a feature vs. its normalized importance in the respective model. Color-coded dashed lines represent the mean R-squared for a model’s features’ age correlations. The probes with the highest importances for all models had |*r*| *<* 0.45. **b** R-squared of the residuals of a feature’s age correlation vs. its normalized importance in the respective model. **c** Negative log of a feature’s p-value (*H*_0_ := no difference in distributions of beta values between controls and samples with inflammaging) vs. its normalized importance in the respective model.

Next, following up on our discovery that DNAm noise is a biomarker of aging [23] we explored the relationship between a selected CpG’s increase in variance with age and its relative importance (**Figure 1b**). To measure age-dependent variance we took the residuals from the age correlations for each CpG and regressed them on chronological age. We then plotted the R-squared value for this regression against the CpG’s relative importance. Once again, we found a negative correlation between age-specific CpG dysregulation and its importance in the EN model: features with the highest noise (|*r*|*≤* 0.4) had the lowest importances, while the CpGs with the highest importance for each model tended to have low noise (|*r*|*≤* 0.24). This suggests that age-related noise is not the primary criteria for feature importance in EN clocks.

Next, we explored the possibility that the elastic net selects CpGs, which are significantly different between healthy people and patients with diseases associated with inflammaging (**Figure 1c**). To do this, we performed a two-tailed Mann-Whitney U test (methods) on the beta value distributions of each CpG probe between healthy controls and patients. We then plotted the negative log of the p-value against the relative importance of each CpG in its respective model. Higher negative log values indicate larger differences between patients and healthy controls for that CpG probe. Again, as with the lack of CpG relevance to age-specific changes in values, or dysregulation, there was a general trend of sharply increasing EN-assigned importances of CpGs that differ *the least* between healthy people and patients, indicating that the elastic net does not select CpGs based on their differential methylation due to inflammaging.

Thus, all examined clocks not only fail to weight DNAm biomarkers of aging by significance but also appear to preferentially exclude such biomarkers. The exclusion of biologically relevant CpGs results from the combination of the L1 penalty for feature selection and regularization, as well as minimizing the EN cost function. This function intentionally penalizes CpGs with little-to-no overall individual correlation with age or its proxies, even if they may correlate differently between various training cohorts (e.g., healthy control versus patient samples), by reducing their weights to zero (**Box 1**). The larger the signal shift between two cohorts in the training data at a CpG probe, the more likely EN is to assign a lower weight to that CpG. However, CpG sites with high degrees of differential methylation are often of the greatest biological interest, as they may indicate key differences related to disease and, consequently, biological age acceleration.

#### Box 1

To illustrate this we first consider the EN cost function

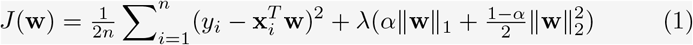

where *n* is number of training samples, *y*_*i*_ is the label (e.g. age) for the *i*th sample, 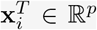 is the sample vector for the *i*th training sample, **w** *∈* R^*p*^ is the vector of weights, and *λ* and *α* are hyperparameters for the regularization penalty and mixing between the L1 an L2 norms respectively.

The weights are learned via gradient descent with the update rule

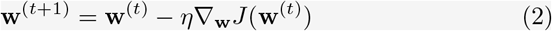

where *t* is the training iteration step, *η* is the learning rate and where

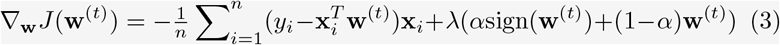

where sign (*·*) is the element-wise sign function. An individual weight 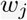is then updated by

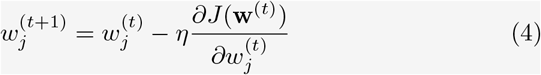

where the gradient component corresponding to the feature weight 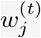 is

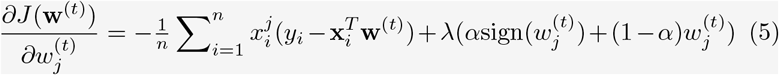

Note that 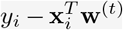 is the *i*th residual 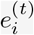 at iteration *t*. Thus, the size of the gradient component corresponding to 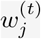 depends in part on the inner product of the *j*th *feature* vector **x**^*j*^ and the vector of residuals **e**^(*t*)^ i.e.

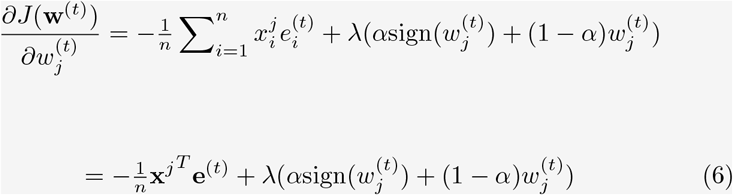

As **w**^(*t*+1)^ converges to the solution ŵ, each 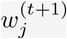 converges to 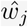, which individually occurs as 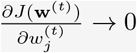. At this limit

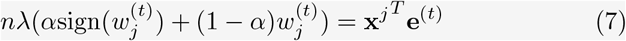

and thus the convergence of the weight 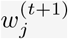 depends on 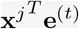, which is a surrogate of the correlation between the *j*th feature and the training residuals at iteration *t*. Note that

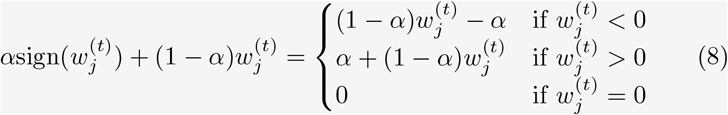

Importantly, *α* is the hyperparameter controlling the balance between the L1 and L2 penalties and thus 0 *≤ α ≤* 1. Under this constraint, it can be observed that 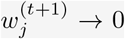 only when 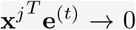, indicating the *j*th feature and training residuals at iteration *t* are uncorrelated. This is the expected outcome under the influence of the L1 penalty, which aims to nullify features lacking correlation with the response variable. However, in the context of DNAm clocks, a deeper examination is warranted regarding the effect which differential methylation could have on the learned weight for a CpG. Given the biological significance of differentially methylated CpGs, their inclusion in any feature subset selection is desirable when constructing models which putatively detect aberrant biological aging.

First, consider the expectation of **e**^*T*^ **x**^*j*^ of the final model where each *e*_*i*_ and 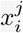 are i.i.d.:

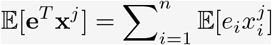

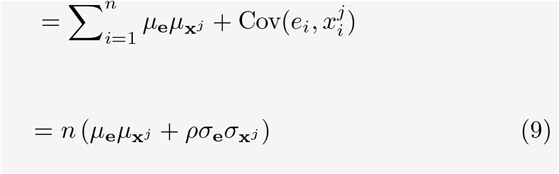

where *ρ* is the correlation between the *j*th feature and the training residuals. However, note that for EN models where *λ* is small the training residuals converge to a normal distribution with mean zero and standard deviation *σ*_**e**_ due to the least squares constraint [31].

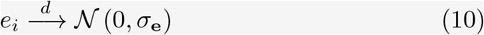

assuming a linear relationship between the features with the response variable, homoscedasticity of the residuals and independence of the residuals–assumptions valid for DNAm clocks and many CpG features. Thus, in the final model *μ*_**e**_ *≈* 0 and

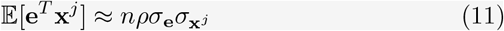

Next, consider a feature **x**^*j*^ that has a non-zero correlation with the response variable, and which is also differentially methylated between two cohorts *A* and *B* within the training set (i.e. 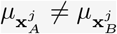). Then

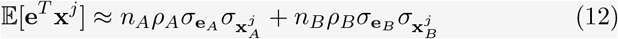

It is therefore feasible to have e^*T*^ **x**^*j*^ = 0, and by consequence *w*_*j*_ = 0, for a feature which *does* correlate with the response variable if

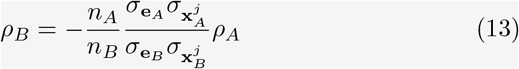

In the context of DNAm clock models, differential methylation at the *j*th CpG would induce a difference in the means of the predicted ages for the two cohorts. The best-fit line between the predicted and actual ages at iteration *t* would be induced to act as a quasi-decision boundary between the two cohorts as the mean of the residuals is pushed towards zero by the least squares penalty (**eqn. (10)**). In such a case the correlations between the beta values and residuals for the two cohorts will be approximately equal and opposite in sign, consistent with **eqn. (13)**. This demonstrates how features with differential methylation between two cohorts within a training set (e.g. a healthy control group and a group with age-associated disease) *could* be excluded from an EN model.

To test this conjecture, we trained several EN models on a dataset consisting of a cohort of rheumatoid arthritis (RA) patients and a healthy control (HC) cohort (GSE42861). We experimented with various mixtures of L1 and L2 penalties and the degree of regularization, using a training feature set conditioned on individual correlation with age. We plotted the learned feature weights against the absolute value of the effect size of the shift between the cohorts’ individual regression lines for the given CpG and age (**Supplementary figure 1a, b**).

The results confirm our conjecture that the L1 penalty, responsible for feature selection in EN models, selects *against* CpGs which are differentially methylated between training cohorts, as CpGs with larger mean shifts between the RA and HC corhorts tended to be assigned smaller weights. Moreover, this trend strengthened with increasing regularization (e.g. higher *λ*). We further confirmed this result for another dataset (GSE72774) consisting of a cohort of Parkinson’s disease patients and a healthy control cohort (**Supplementary figure 1c**). Together, the mathematical rationale and empirical results demonstrate that the larger the signal shift between two cohorts in the training data at a CpG probe, the more likely it is for EN to assign a lower weight to that CpG.

### Quantifying Normal Variance in DNAm Clock Residuals

One method for predicting biological age with DNAm clocks is to use the residual, or difference between sample’s predicted age and their known chronological age, with deviations suggested to indicate age acceleration [1, 3–5, 7–12]. However, residuals would be expected to not only include biological age acceleration, but also natural variation in the healthy human methylome as well as random experimental variation during the assays, data processing, and variance due to approximating a non-linear process with a linear model.

To quantify normal variation and determine thresholds for potentially aberrant methylomes, we fit a probability distribution to the residuals of predicted ages for healthy controls from the composite dataset (GSE42861, GSE125105, GSE72774, GSE106648) for each predictive model. The residuals were approximately normally distributed according to Q-Q plots (**Supplementary figure2a**). Thus, a normal distribution was fit to the residuals for each model (**Figure 2a**). We also investigated the residual distributions at ten-year age intervals for each model (**Supplementary figure 2b**).

**Fig 2:**
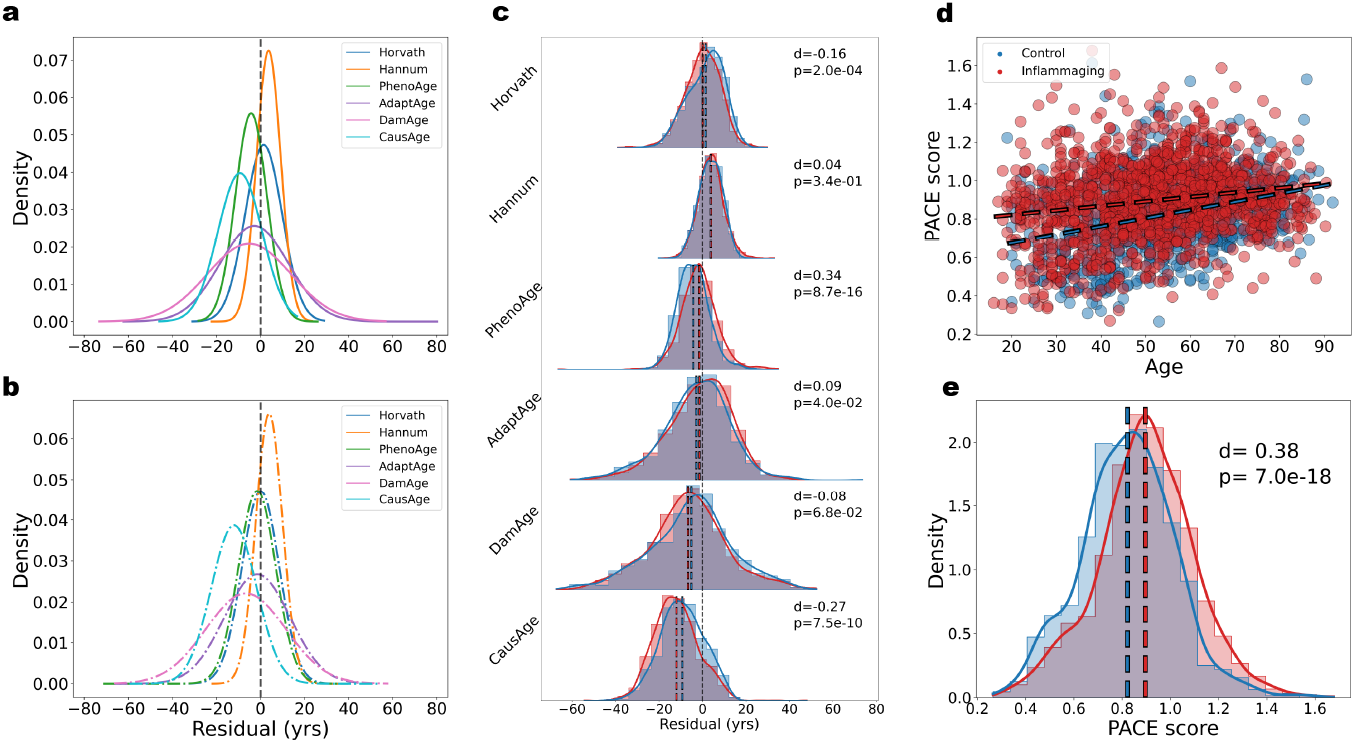
**a** Fitted normal distributions for the residuals of first- and next-generation clocks on healthy controls in the composite dataset. **b** Fitted normal distributions for the residuals of clocks on the inflammaging cohort in the composite dataset. **c** KDE plots with histograms normalized by cohort size comparing the distributions of residuals for healthy control cohort (blue) and inflammaging cohort (red) in the composite dataset for first- and next-generation clocks (d := Cohen’s d). **d** DunedinPACE score calculated for the healthy control and inflammaging cohorts vs. chronological age. Blue and red dashed best-fit lines are for the healthy and inflammaging cohorts respectively. **e** Normalized KDE plots and histograms comparing the distributions of PACE scores for the healthy and inflammaging cohorts. Corresponding dashed lines represent cohort means.

Next, we calculated the 95% interpercentile range (95-IPR, i.e., the range containing the central 95% of residuals) for the healthy cohort (HC) for each model. Additionally, we calculated the means of the residuals of the healthy cohort and inflammaging cohort (IC) to quantify how well-calibrated each model is to test data. We define good calibration as having a residual mean close to zero, indicating there is no bias for having more positive or more negative residuals. These results, summarized in **Table 1**, demonstrate that, when correcting for mean shift, residuals of up to 10.8 years fall within the expected range for healthy individuals for the model with the smallest 95-IPR and thus highest precision (Hannum). Averaging over all models tested, a residual of over 24.6 years would be needed to be outside the expected range for healthy individuals, when correcting for mean shifts. The mean shifts for the inflammaging cohort for all models is less than this threshold by an order of magnitude, suggesting that all models lack sufficient power to detect the age accelerating effects of inflammaging (**Table 1, Figure 2b**).

**Table 1:** Residual Distribution and Calibration of DNAm Clocks.

| Model | HC: 95-IPR range (yrs) | HC: mean (yrs) | IC: mean shift (yrs) |
| --- | --- | --- | --- |
| AdaptAge | -33.4 – 27.7 | -2.8 | 1.3 |
| DamAge | -42.6 – 32.3 | -5.2 | -1.4 |
| CausAge | -29.0 – 10.4 | -9.3 | -2.7 |
| Hannum | -7.1 – 14.5 | 3.7 | 0.2 |
| Horvath | -15.1 – 18.0 | 1.4 | -1.3 |
| PhenoAge | -18.3 – 9.8 | -4.3 | 2.8 |

We performed a two-tailed Welch’s t-test to determine statistical significance between the distributions of residuals for the healthy and inflammaging cohorts (**Figure 2c**). Importantly, high numbers of samples can yield artificially low p-values so we also quantified effect size, which is more indicative of the power of a model. To this end we used Cohen’s d, which is a signal-to-noise ratio, and we use the effect size rule of thumb suggested by Sawilowsky (2009) [32]. Of the models, only PhenoAge and AdaptAge had statistically *higher* means of the residuals (*p <* 0.05) for patients, but both have small effect sizes. Moreover, the higher mean shifts for the inflammaging cohort were well within the 95-IPR for both models. These results suggest that all the models investigated lack sufficient power to detect the age accelerating effects of inflammaging via residual.

Next, we analyzed the DunedinPACE model separately as it yields a score rather than a predicted age, with a higher score indicating a higher rate, or pace of aging [6]. We first generated a PACE score (methods) for each sample in both the healthy and patient cohorts of the composite dataset and plotted the PACE score versus chronological age for both (**Figure 2d**). PACE was trained on samples from people who were all *∼*53yrs, yet interestingly, it worked as an age predictor when tested on the combined dataset that spans 20-90+yrs. This observation demonstrates that training predictors that are proxies of chronology, e.g., instances of disease increase with aging, continue to predict age.

Next, to determine if PACE detects inflammaging we performed a two-tailed Welch’s t-test, comparing the means of the distributions between the cohorts and calculated the effect size (**Figure 2e**). While the mean of the inflammaging population slightly shifted upwards with *d* = 0.38, this is a small effect size for inflammaging, a hallmark of aging that should be robustly discernable.

Summarily, most models failed in distinguishing health from disease and even PACE, which was trained on extreme differences between people who were healthy, vs. those who had heart attacks, strokes, etc., had only a marginal power to detect the age-accelerating effects of inflammaging. Overall, there remains room for improvement in feature selection.

### OLS regression suffices for a generalizable age predictor with hypothesis-driven feature selection

DNAm clocks trained on DNA methylation array datasets typically have tens-to-hundreds of thousands of CpG features (*p*) and hundreds to a few thousand samples (*n*). Ordinary least squares regression has the advantage of good interpretability on the correlations between the response and predictor variables, but on this structure of data, where *p >> n*, OLS tends to have significant overfitting issues and has very high variance on out-of-sample data [2]. Elastic net regression models combat this issue via regularization, which does feature selection and introduces bias to the model, effectively reducing variance on out-of-sample data. This makes EN more generalizable than ordinary least squares models. However, as demonstrated, EN clock models suffer a lack of biological interpretability and may select against features with the most biological significance.

We hypothesized that, due to the presence of many CpGs with high linear age correlations, it is possible to construct a generalizable and interpretable simple ordinary least squares age predictor from a reduced set of CpGs that have a priori biological relevance. To test this hypothesis, we implemented a modified version of the forward stepwise selection (mFSS) algorithm (**Figure 3a, Algorithm 1**).

**Fig 3:**
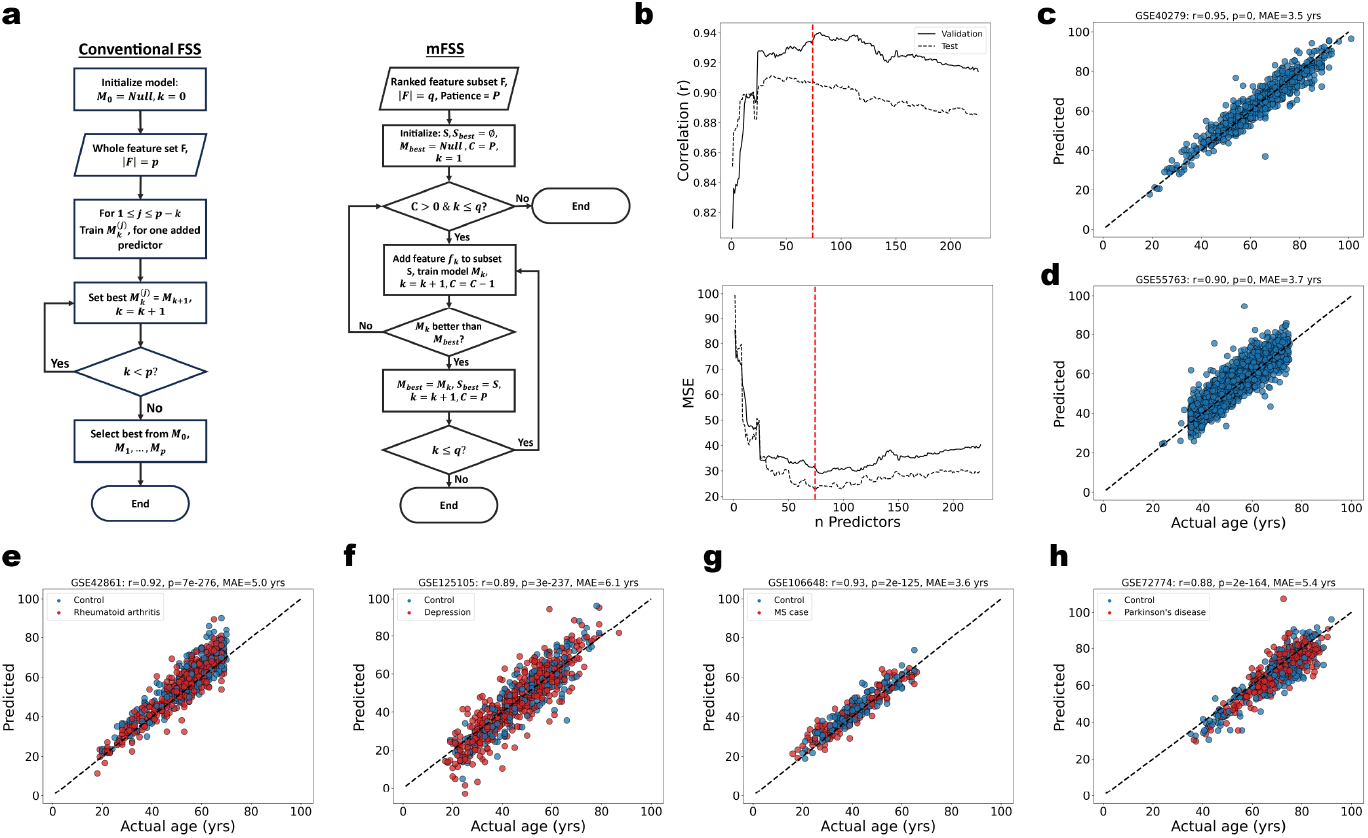
**a** Schema of the the conventional forward stepwise selection algorithm versus the modified forward stepwise selection implementation. **b** (Top) Pearson correlation vs. number of features included in model. (Bottom) Mean squared error vs. number of features included in model. Vertical dashed red line indicates optimum number of features. **c** The optimum mFSS model predictions on the training set (GSE40279). Black dashed line is y=x. **d** mFSS model predictions on the test set (GSE55763) **e** mFSS model predictions on out-of-sample test data with rheumatoid arthritis patients and healthy controls (GSE42861) **f** mFSS model predictions on out-of-sample test data with depression patients and healthy controls (GSE125105) **g** mFSS model predictions on out-of-sample test data with multiple sclerosis patients and healthy controls (GSE106648) **h** mFSS model predictions on out-of-sample test data with Parkinson’s disease patients and healthy controls (GSE72774).

Briefly, we begin with a list of features, ranked by some biological criterion (e.g. correlation with age, increase in noise with age etc.). An OLS regression model is trained as each feature is sequentially added to the model, recording the value of a cost function (e.g. mean squared error) on a test set at each iteration. If the addition of a feature achieves a new low for the cost function, we record that model as the best model. We employ a patience hyperparameter *P* which allows for up to *P* unproductive feature additions (i.e. producing a model with a new minimum cost) before stopping and returning the last best model with its optimal feature selection. The algorithm stops either after *P* unproductive additions or after all features in the initial list have been added. We generated a ranked list of features from a 450K DNAme array dataset (GSE40279, N=656) by regressing chronological age on each CpG feature independently, ranking the CpGs by the magnitude of their age correlation. This same dataset was used for training, and we used an out-of-sample dataset (GSE55763, N=2,631) as a test set. We next trained an mFSS OLS model, splitting the training data (85/15) into training and validation sets (**Figure 3a, b**). The selected features and weights for this model is in **Supplementary Table 1**. The top ranked CpG alone yields a model with *r >* 0.8, and the top 74 CpGs yielded the best OLS model with *r* = 0.9 and mean absolute error (MAE) of 3.7 years in the test set (**Figure 3b-d**). The mFSS clock displays high accuracy and low variance on out-of-sample data, demonstrating that it is generalizable and does not suffer from overfitting, **Figure 3e-h**. Importantly, the mFSS clock trained on CpGs with the highest age correlation is well-calibrated to test data, out-performing published EN-trained clocks.

The biological relevance of the genes annotated to the CpGs used in our mFSS OLS model is known and their inclusion in the model is due to their age-specific change in DNA methylation. An analysis of the roles of the top CpG annotated genes of our mFSS that overlap between the 450K datasets, suggests that they mostly reflect the aging-related changes, such as increase in body mass/fat, repeated instances of viral infections and inflammation, pre-cancerous growths, alcohol detoxification, changes in teeth/hair/skin, etc., **Supplementary Table 2**. Other notable roles include protein degradation, maturation of the extracellular matrix and perception of pain.

### Coherence theory of disease-related shifts for linear models

It has been suggested, without empirical verification by randomized blinded studies, that deviations between predicted and actual ages signify age acceleration [1, 3–5, 7–12], yet our data shows that such deviations could be also due to random experimental variation and healthy dynamics of DNAm. Additionally, EN clocks do not reliably distinguish healthy people from patients diagnosed with diseases, including those of inflammaging ([23] and **Figure 2**). At the same time, as expected, there are some CpGs that reliably detect disease-specific methylation; **Supplementary figure 3a** shows two such cases for rheumatoid arthritis. In solving this conundrum where disease vs. health definition is low in linear models, but DNAme changes with disease are pronounced in the DNAme array data, we note that both increased and decreased shifts in DNAme that are disease-specific are possible, and pairing those signal shifts with weights, which can be either positive negative, in linear combination may cause a negation of the disease signal present at individual CpGs. It would be more robust and better reflect the biology to pair a positive shift with positive weight, and negative with negative.

In linear models, the signs of the weights are determined by the solution which minimizes the cost function of the model and need not depend on the signs of the correlations between individual features and the response variable (i.e. a feature with positive association with the response variable may still be assigned a negative weight in the multiple regression solution, and vice versa). Similarly, individual feature shifts can be either positive or negative, depending on the agreement in sign between the model weights and signal shifts (**Box 2**). This can cause an artificial cancellation effect and can produce a net age shift near or equal to zero despite significant differences between the methylomes of disease and healthy states. We define such a property of a linear model as model *incoherence*.

#### Box 2

We investigate the potential negation effect by considering a linear model (e.g. EN or mFSS OLS) with weights *w*_*i*_ for 1 *≤ i ≤ p*. A selected CpG probe subset of size *p*, yields an age prediction *ŷ* via a linear combination of the model weights and a single sample’s beta values for the selected CpGs.

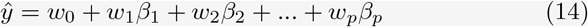

We define a hypothetical disease-related signal shift at a particular CpG probe as the difference of beta values *β*_*d*_ and *β*_*h*_ between the healthy and disease states respectively:

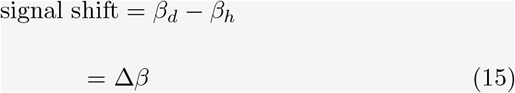

We similarly define a hypothetical disease-related shift in age prediction for a given linear model as the difference in the age predictions, *ŷ*_*d*_ and *ŷ*_*h*_ between the disease and healthy states respectively:

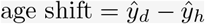

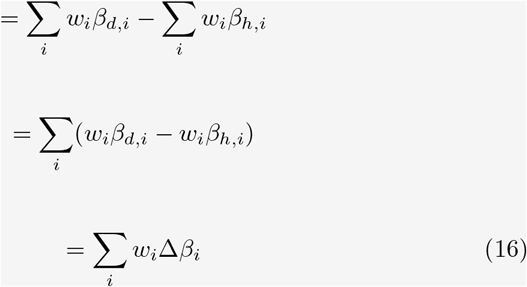

Therefore, a hypothetical disease-related shift in age prediction is due to the sum of weighted signal shifts. We define each *w*_*i*_Δ*β*_*i*_ as the *feature shift* between the disease and healthy states for a given linear model. Therefore a hypothetical age shift associated with disease is the sum of individual feature shifts:

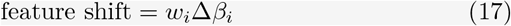

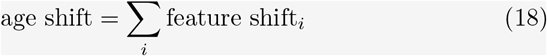

De-coherent feature shifts where the signal shift for a given disease is negative, can be eliminated by simple reflection in the training and test data: 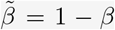, such that if Δ*β ≤* 0, then 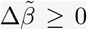. **Supplementary figure 3c** demonstrates the effect of this transformation on toy data. These two modifications to the training criteria and input data taken together constitute feature rectification and it is summarized as

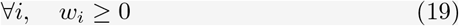

and

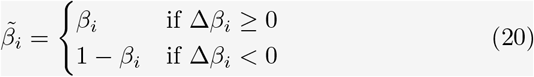

To demonstrate the principle of feature rectification, we created a synthetic, or ‘toy,’ dataset comprising separate cohorts with signal shifts present in certain features. **Supplementary figure 3d** contrasts linear models trained on the toy data with and without feature rectification and its effect on model coherence. Without feature rectification, a linear model is simply trained on putative healthy training data and there is a net cancellation of coherent and de-coherent feature shifts when predicting on the test data containing a condition cohort, producing a linear model which does not yield resolution between the condition and healthy cohorts. When feature rectification is implemented, the signal shifts of the condition cohort are first calculated (methods) and the reflection transformation is applied to both the training and test data. A new model is then trained with the model weights restricted to be positive. The coherent feature shifts in the test data yield an overall coherent model where the condition cohort is shifted upward relative to the healthy control cohort, providing resolution between them and effectively establishing separate aging trajectories for individuals with and without the condition.

Feature rectification is a condition-dependent procedure, as it requires a priori knowledge of the signal shifts for the particular condition. Additionally, a model trained on feature-rectified data will only be valid for test data which has also undergone the reflection transformation on the probes which are known to have negative shifts for the condition under investigation.

Conversely, we define the property where all feature shifts are in agreement in sign to produce a net positive age shift as model *coherence*. Due to the association of *overprediction* of age with age acceleration, we define positive feature shifts as *coherent* and negative feature shifts as *de-coherent*. The degree of incoherence in a model is expected to vary for different diseases and to depend on the interactions between the feature shifts for a given disease. **Supplementary figure 3b** summarizes the interactions between the signal shifts and model weights which produce coherent and de-coherent feature shifts.

Model coherence is a desirable property if researchers would like their DNAm age predictors to not only predict chronological age, but to also detect shifted trajectories of biological age. Here, we propose a method, which we term *feature rectification*, for inducing model coherence during model training by allowing only positive feature shifts. While the signs of feature weights for a model cannot be known a priori, it is possible to constrain a model to have non-negative weights, thereby eliminating de-coherent feature shifts, where signal shifts are positive.

### mFSS OLS for coherent, generalizable age predictor and inflammaging detector

Following the success with the toy data, we tested our coherence theory for linear models by training a mFSS model and with feature rectification for actual PBMC samples from people with rheumatoid arthritis (RA) and healthy controls. We calculated the RA signal shifts in the GSE42861 dataset (**Supplementary Table 3**) for the top 10,000 age correlated CpGs (with respect to the samples in GSE40279) and performed the reflection transformation for the negatively shifted probe signals in the RA test set and the GSE40279 and GSE55763 training and validation datasets respectively (methods). Next, we trained a mFSS OLS model constrained to have non-negative weights. The features were fed into the model ranked by the magnitude of their age correlation in the training set. The validation set MSE was optimized at the 680th feature, with 124 features having non-zero weights (selected features and weights, and feature rectification information in **Supplementary Table 4**, training and validation performance in **Supplementary figure 4a**).

We next obtained age predictions for the RA test set and plotted them against the actual ages. **Figure 4a** compares the incoherent, basic mFSS OLS model predictions and the mFSS OLS model trained on rectified features. The coherence is apparent for the latter model, with the RA cohort predictions visibly shifted upward relative to the healthy control (HC) cohort. We quantified the shift by performing a two-tailed Welch’s t-test on the distributions of the residuals of the HC and RA cohorts (*p* = 1.9*e −*25) and calculating the effect size of the shift (*d* = 0.83), **Figure 4b,c**. Although the RA dataset was not used to train the model directly, the feature rectification was based on the signal shifts between the RA and HC cohorts in the GSE42861 dataset. To test whether the coherence effect is still present in completely out-of-sample data we predicted the ages on several other datasets with different disease cohorts (multiple sclerosis, depression, Parkinson’s disease and tauopathies), **Supplementary figure 4b-e**. Each of the disease cohorts was slightly up-shifted relative to their respective healthy control cohorts, but with very small effect sizes ranging from *d* = 0.14 to *d* = 0.19.

**Fig 4:**
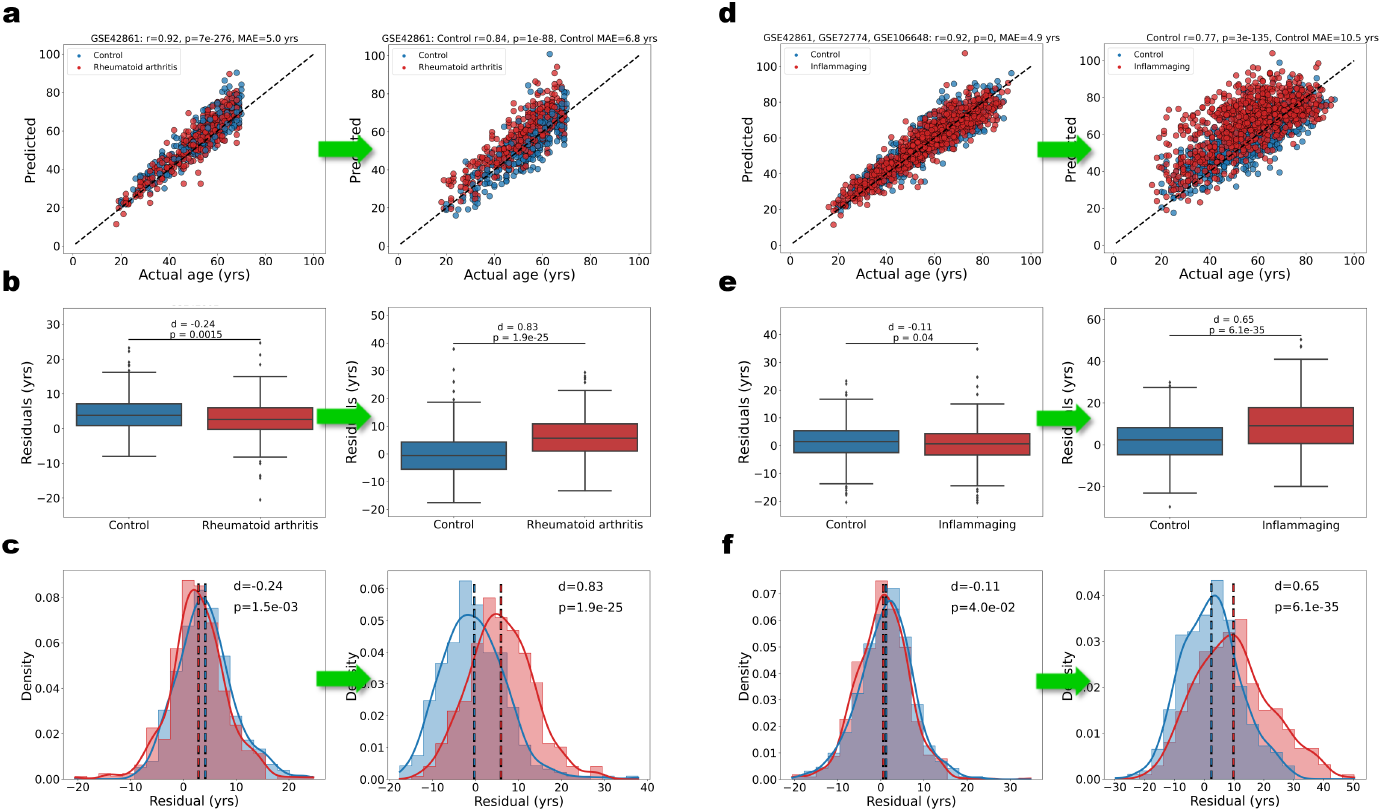
**a** (Left) Scatterplot of ages predicted by the basic mFSS clock vs. actual ages for the GSE42861 dataset colored by disease state (healthy controls in blue; rheumatoid arthritis in red) compared to predictions of a coherent mFSS model trained on rectified features based on RA signal shifts (right). Correlation, p-value and MAE calculated for the control cohort only. **b** Box plots of the residuals for healthy controls and rheumatoid arthritis patients corresponding to the basic mFSS model (left) and coherent mFSS model trained on rectified features (right). p-values determined by Welch’s t-test. Effect size (d) determined by Cohen’s d (methods). **c** Histograms and KDE plots, normalized by cohort size, of the residuals for healthy controls and rheumatoid arthritis patients corresponding to the basic mFSS model (left) and coherent mFSS model trained on rectified features (right). Vertical lines indicate the cohort means. **d** Scatter plot of predicted age vs. actual age for a combined dataset (GSE42861, GSE72774, GSE106648) colored by disease state (healthy controls in blue; inflammaging-associated disease in red), compared to predictions of a coherent mFSS model trained on rectified features based on inflammaging signal shifts (right). Correlation, p-value and MAE calculated for the control cohort only. **e** Box plots of the residuals for corresponding to the basic mFSS model (left) and coherent mFSS model (IR-mFSS model) trained on rectified features (right) on the combined dataset. **f** Histograms and KDE plots, normalized by cohort size, of the residuals for the basic mFSS model (left) and coherent mFSS model (IR-mFSS model) trained on rectified features (right) on the combined dataset. Vertical lines indicate the cohort means.

Interestingly, training on features ranked instead by the magnitude of the *signal shifts* between healthy controls and RA reduced the accuracy of predictions of chronological age in all datasets and decreased calibration, while increasing the resolution between health and disease for all but one disease (Parkinson’s) (**Supplementary figure 5b-f**). The MSE for the validation set was optimized at the 1,656th feature with 47 features having non-zero weights (**Supplementary figure 5a**).This is expected, because to linearly predict age with lowest error, the CpGs that change the most with disease between people of the *same age* will be minimized via the least squares constraint.

Next, to make a more generalized disease coherent model we trained a mFSS OLS model with feature rectification based on a combined dataset with inflammaging-associated disease cohorts (GSE42861, GSE106648, GSE72774), which we dubbed the inflammaging-rectified mFSS (IR-mFSS) model. The model was again trained on GSE40279 and validated on GSE55763, with the top 10,000 age correlated CpGs (per the GSE40279 samples) ranked by absolute signal shift magnitude (**Supplementary Table 5**) for the combined inflammaging dataset (selected features and weights, and feature rectification information in **Supplementary Table 6**, training and validation performance in **Supplementary figure 6a**). **Figure 4d** compares the incoherent basic mFSS OLS model predictions on the combined inflammaging dataset and the coherent IR-mFSS model. We quantified the shift by performing a two-tailed Welch’s t-test on the distributions of the residuals of the HC and inflammaging cohorts (*p* = 6.1*e −*35) and calculating the effect size, which was medium-to-large (*d* = 0.65), as seen in **Figure 4e**,**f. Supplementary figure 6b-f** shows the distributions of the residuals for different datasets, comparing patients with inflammaging-associated diseases and healthy controls. The IR-mFSS clock increased the resolution over the basic mFSS clock and substantially for all diseases, which effect sizes ranging from small (d=0.27) to very large (d=1.3).

The genes annotated to the 47 CpGs selected by the IR-mFSS clock and their functions are provided in (**Supplementary Table 7**).

## Discussion

This study establishes why linear machine learning models, such as EN clocks, underperform in their ability to distinguish health from many diseases, including inflammaging. Specifically, we show that linear EN regression models, whether trained to predict age or health scores, or mortality, etc., all the major models trained on DNA methylation have feature selections without clear biological meaning. The features selected may simply minimize the EN cost function. Our results show that CpG importance in the DNAm clocks is assigned in disregard of the age-specific changes in DNAm; it remains to be seen what biological process, if any, plays a role in the assigned weights, or if the subsets are simply a mathematical artifact of model optimization and batch effect.

We also determined that many of the EN-trained models are poorly calibrated for out-of-sample data and have very large natural variance in prediction errors for healthy people. Thus, there is no reason to think about loss of health or age acceleration unless a residual is more than 10-20 years, and until a blinded randomized study empirically verifies the predictive accuracy of a given model.

To improve the biological relevance of DNAm age predictors, we developed a modified forward stepwise selection OLS regression model with a clear hypothesis-driven feature selection. We demonstrated that such mFSS models are ML independent yet are generalizable and better calibrated than EN clocks when trained on features with at least some correlation with age.

Importantly, we identified a key detriment of linear models when using residuals as a measure of age acceleration, i.e., an incoherence between combinations of multiple CpG features, some of which shift in their biological DNAm in an opposite direction of the model-assigned weights. Our theory for linear model coherence explains why linear models tend not to provide resolution between health and disease despite significant disease-imposed changes in CpG beta values. This theory is confirmed by the demonstration that feature rectification yields coherent, generalizable linear models through mFSS OLS, which can detect aging and simultaneously, disease. Of note, while feature rectification provides a necessary correction to enable linear models to detect inflammaging, neither biological aging, nor most diseases are linear processes, so interpretation of apparent differences in aging trajectories between healthy and patients’ cohorts should be made with caution.

## Methods

Pre-processing of raw IDAT files (when available) was performed in R version 4.3.1. All other data handling was performed in Python version 3.8.16. All statistical analysis was performed with the *stats* module in the SciPy library, version 1.10.0. All linear modeling was performed with the relevant modules in the scikit-learn library, version 1.2.0.

### Dataset pre-processing

Raw IDATs were pre-processed using the Minfi library. Noob pre-processing was applied to the entire dataset first and then BMIQ (wateRmelon library) quantile normalization.

Samples which had more than 5% missing values were removed from their respective datasets.

### Imputation of missing values

KNNImputer (with k=2) from the *sklearn*.*impute* module was used to impute missing values.

### Statistical analysis and measure of effect size

Welch’s t-test for unequal variances or the non-parametric Mann-Whitney U test was performed for all hypothesis testing. Cohen’s d with pooled standard deviation was used to measure the effect size between sick cohorts and their corresponding healthy control cohorts:

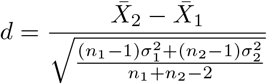

### Analysis of the CpG selection of EN-trained DNAm clocks

All analyses of the CpG selection of first and next-gen clock models was performed on the combined dataset (GSE42861, GSE125105, GSE106648, GSE72774).

#### Age correlation of individual CpG probes

The healthy cohort of the combined dataset was used to measure the age correlation of each CpG probe in the Illumina HumanMethylation450 BeadChip array. The sample beta values for each CpG probe were regressed on the sample ages. The coefficient of determination (R-squared), standard error, and slope and intercept of the best-fit line was recorded for each regression.

#### Normalized feature importance

The normalized feature importance for the *i*th CpG probe in a given model’s CpG probe selection was computed by:

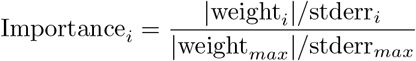

where stderr_*i*_ is the standard error of the regression of the *i*th CpG probe when its beta values are regressed on age.

#### Age-dependent CpG probe variance

The slope and intercept for the best-fit line of a probe’s age correlation gives a sample’s predicted beta value 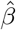 by

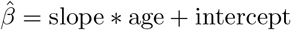

The residual for a sample for a given probe was calculated as

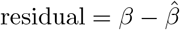

The absolute value of the residuals for a probe were then regressed on age.

#### Disease-associated differential methylation of CpG probes

For each CpG probe the Mann Whitney U-test was performed on the beta values between the healthy control and patient cohorts of the combined dataset, with *H*_0_ being there is no difference between their distributions.

### Generating published model predictions

The age predictions and PACE scores for the combined dataset were calculated by taking the dot product of sample beta values and the model weights provided in the supplementary files for the relevant models.

For the Horvath clock predictions, the inverse of the log transform described in the paper was applied to the dot prodcut to generate the final age predictions.

### Average signed distance between regression lines

The slope *m*_*A*_ and intercept *y*_0,*A*_ for a best-fit line were used to generate

**ŷ**_*A*_ = *m*_*A*_**x** + *y*_0,*A*_**1**_100_, for **x**^*T*^ = (1, 2, …, 100). Next,

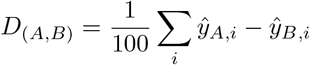

*Note that *D*_(*A,B*)_ = *−D*_(*B,A*)_.

### Effect size between regression lines

The average distance between lines was calculated as described above. This average distance was divided by the pooled standard error of the two regression lines.

### DNAm clock prediction variance

The ages for the healthy control and disease cohorts of the combined dataset were predicted for a given model by taking a linear combination of the beta values for the model’s selected CpG probes and their model weights. Next, the residuals were separately calculated and normality of their distribution was tested by quantile-quantile plot. A normal distribution was then fit to the residuals using the *stats*.*norm*.*fit()* method from the SciPy library.

### Feature rectification for model coherence

#### Calculating signal shifts for a disease

Datasets containing healthy control *H* and patient *D* cohorts were split accordingly. Each CpG was individually regressed on age in the separate cohorts. The average signed distance 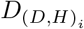 for the *i*th CpG between the best-fit lines between the healthy and patient cohorts was calculated as described above.

#### Reflection transformation

The beta values for the *i*th CpG in the training data for a disease-rectified model were transformed by:

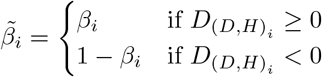

## Data availability

All data used in this study was downloaded from the Gene Expression Omnibus (GEO) repository: https://www.ncbi.nlm.nih.gov/geo/

Raw IDAT files were downloaded and pre-processed (methods) when available. For all others, the processed datasets were downloaded.

**Datasets used:** GSE42861, GSE125105, GSE40279, GSE55763, GSE106648, GSE72774, GSE53740

## Code availability

All relevant code can be accessed at https://github.com/SkinnerCM/conboy-laboratory

## Supplementary Figures

**Supplementary Figure 1.**
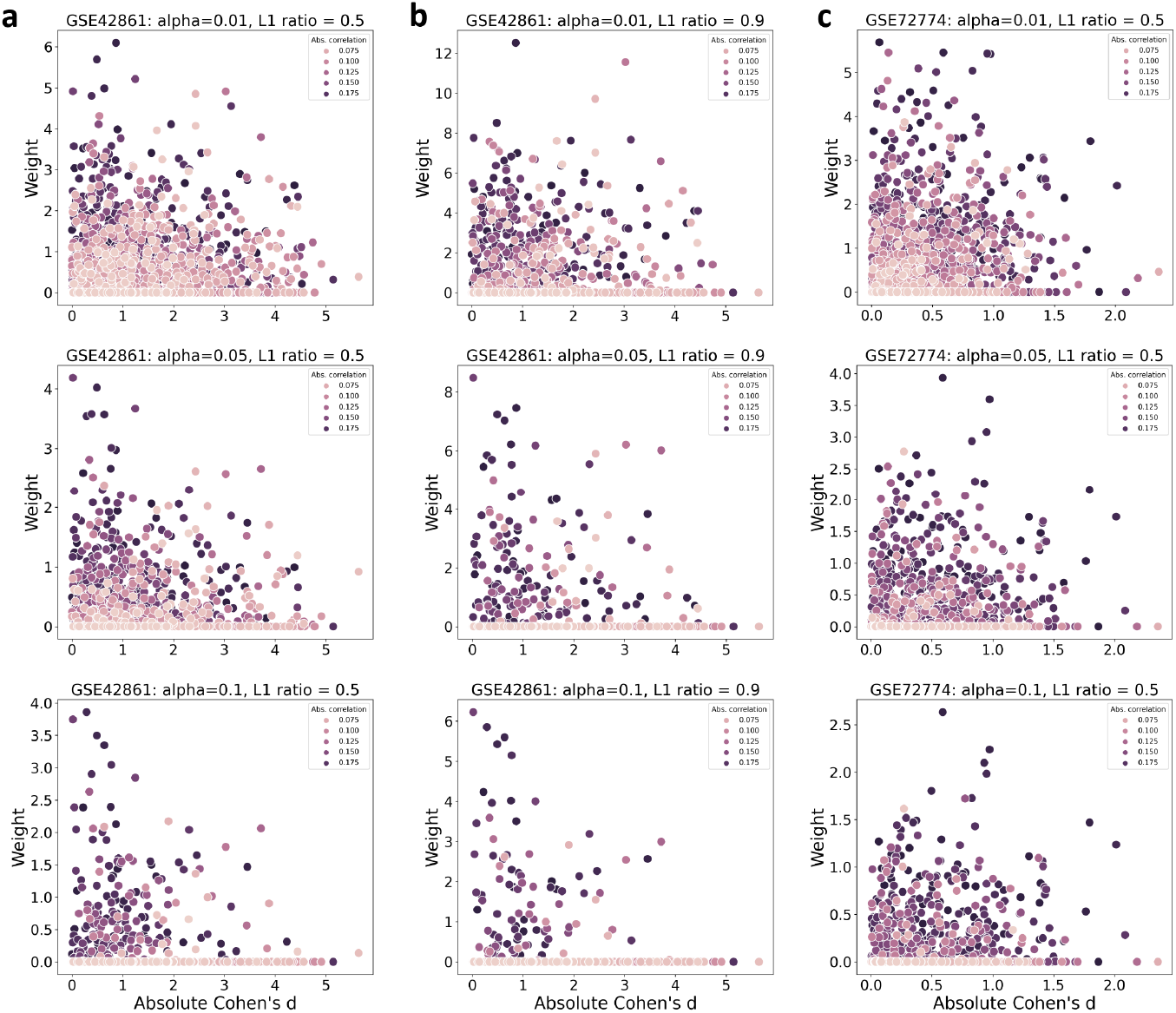
**a** GSE48261: learned EN weight vs. absolute value of effect size (Cohen’s d) with l1 ratio = 0.5 and (top) alpha = 0.01 (middle) alpha = 0.05 (bottom) alpha = 0.1. **b** GSE48261: EN weight vs. absolute value of effect size with l1 ratio = 0.9 and (top) alpha = 0.01 (middle) alpha = 0.05 (bottom) alpha = **c** GSE72774: EN weight vs. absolute value of effect size with l1 ratio = 0.5 and (top) alpha = 0.01 (middle) alpha = 0.05 (bottom) alpha = 0.1.

**Supplementary Figure 2.**
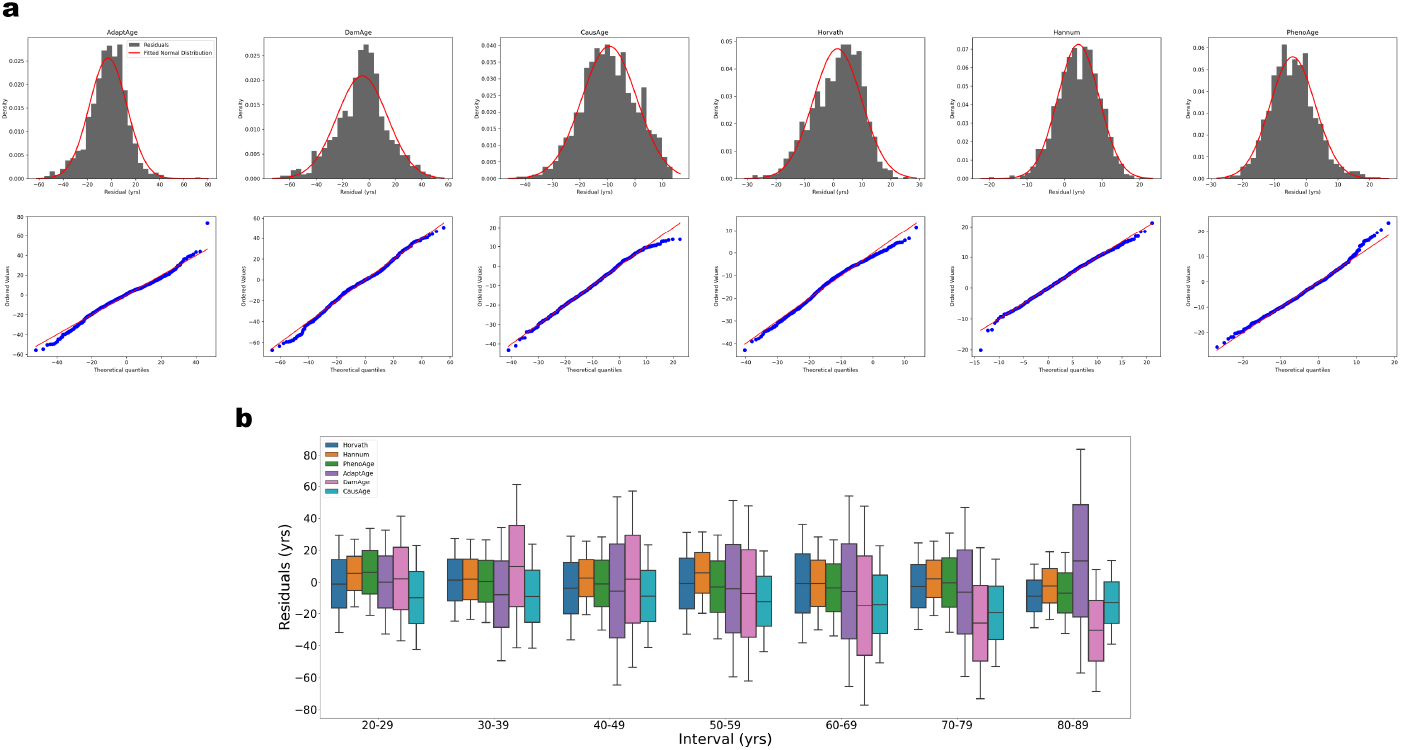
**a** (top) Histograms and fitted normal distributions of the residuals of the age predictions on the healthy cohort of the composite test dataset. (bottom) Q-Q plots for each model showing the goodness-of-fit of the normal distribution to the residuals. **b** 10-year age bins of the residuals for the healthy cohort of the composite test dataset for each of the tested models.

**Supplementary Figure 3.**
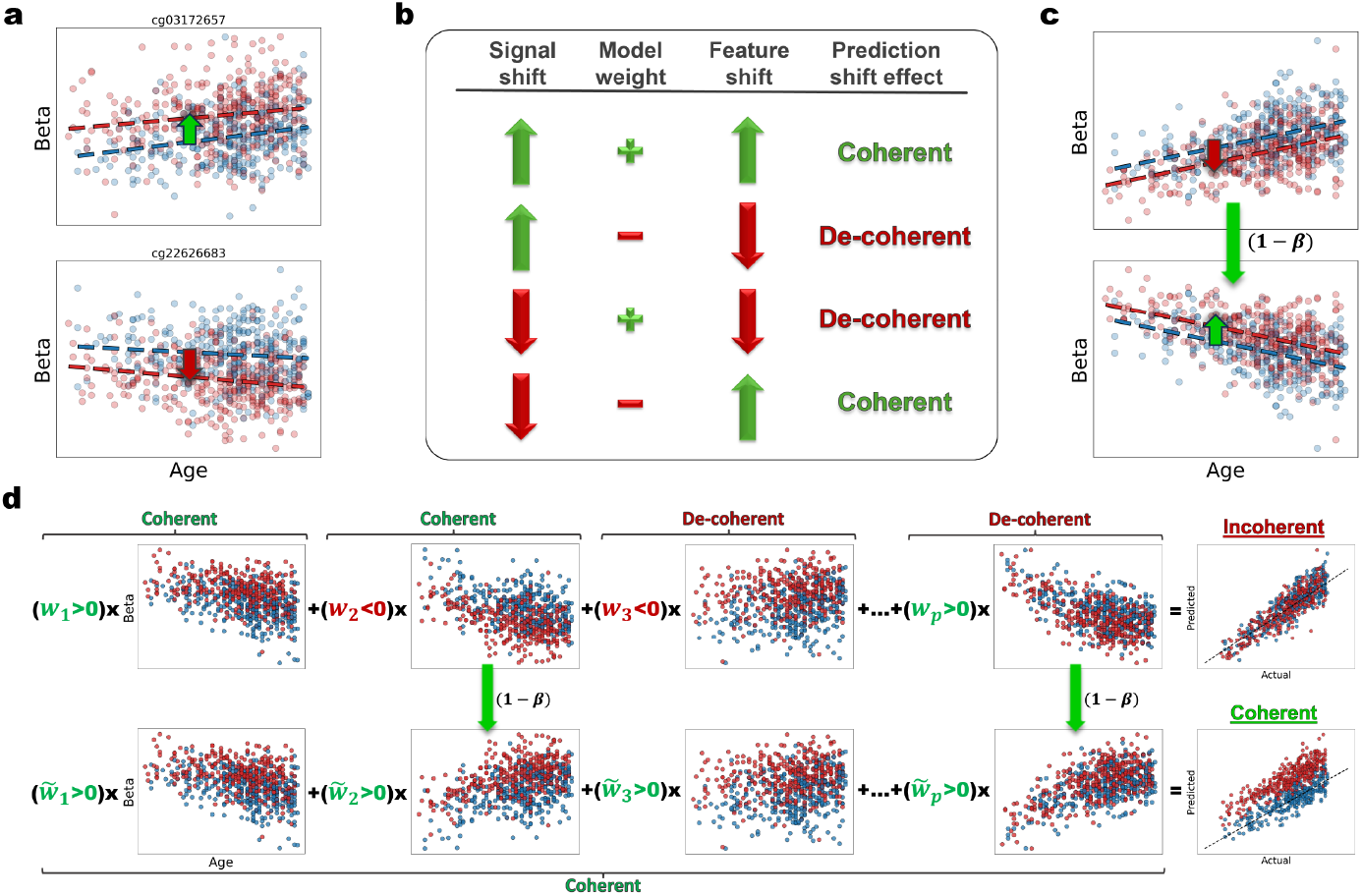
**a** Beta value versus age plotted for healthy controls (blue) and rheumatoid Arthritis patients (red) for representative CpGs in the GSE42861 dataset. (Top) A CpG for which the best-fit line for the RA cohort is shifted up from the HC cohort. (Bottom) A CpG for which the best-fit line for the RA cohort is down-shifted relative to the HC cohort. **b** Table summary of how the product of the weights and disease shifts for model probes combines to produce either coherent or de-coherent feature shifts. **c** Demonstration of the reflection transformation of the beta values for feature shift rectification (toy data). **d** (Top row) Visualization of how the linear combination (weights times beta values) of both coherent and de-coherent feature shifts produces an incoherent picture of biological age-acceleration in the presence of disease. (Bottom) Visualization of how the a linear combination of the rectified feature shifts (positive weights, reflected beta values for CpGs down-shifted for a disease) produce a coherent image of the biological age acceleration due to disease.

**Supplementary figure 4.**
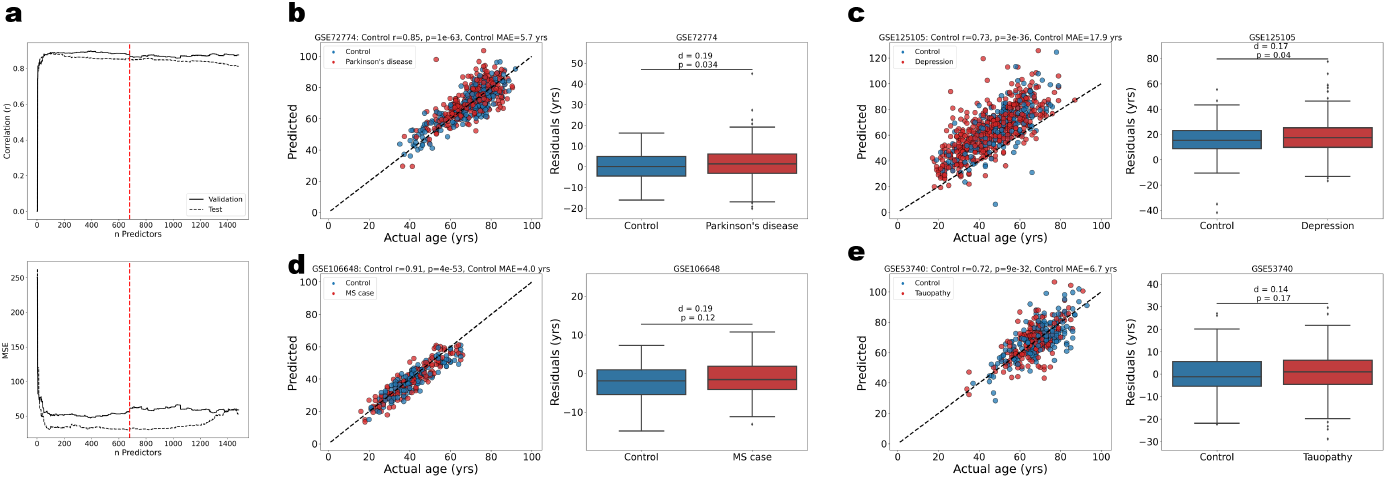
**a** (top) Correlation vs. number of features for RA coherent mFSS model trained on features ranked by age correlation. (bottom) Corresponding plot for MSE vs. number of features. Red dashed line indicates number of features which give the optimum validation MSE. **b** (Left) scatter plot for predicted age vs. actual age for GSE72774. (right) box plot of the residuals of the predictions. **c** (Left) scatter plot for predicted age vs. actual age for GSE125105. (right) box plot of the residuals of the predictions. **d** (Left) scatter plot for predicted age vs. actual age for GSE106648. (right) box plot of the residuals of the predictions. **e** (Left) scatter plot for predicted age vs. actual age for GSE53740. (right) box plot of the residuals of the predictions.

**Supplementary figure 5.**
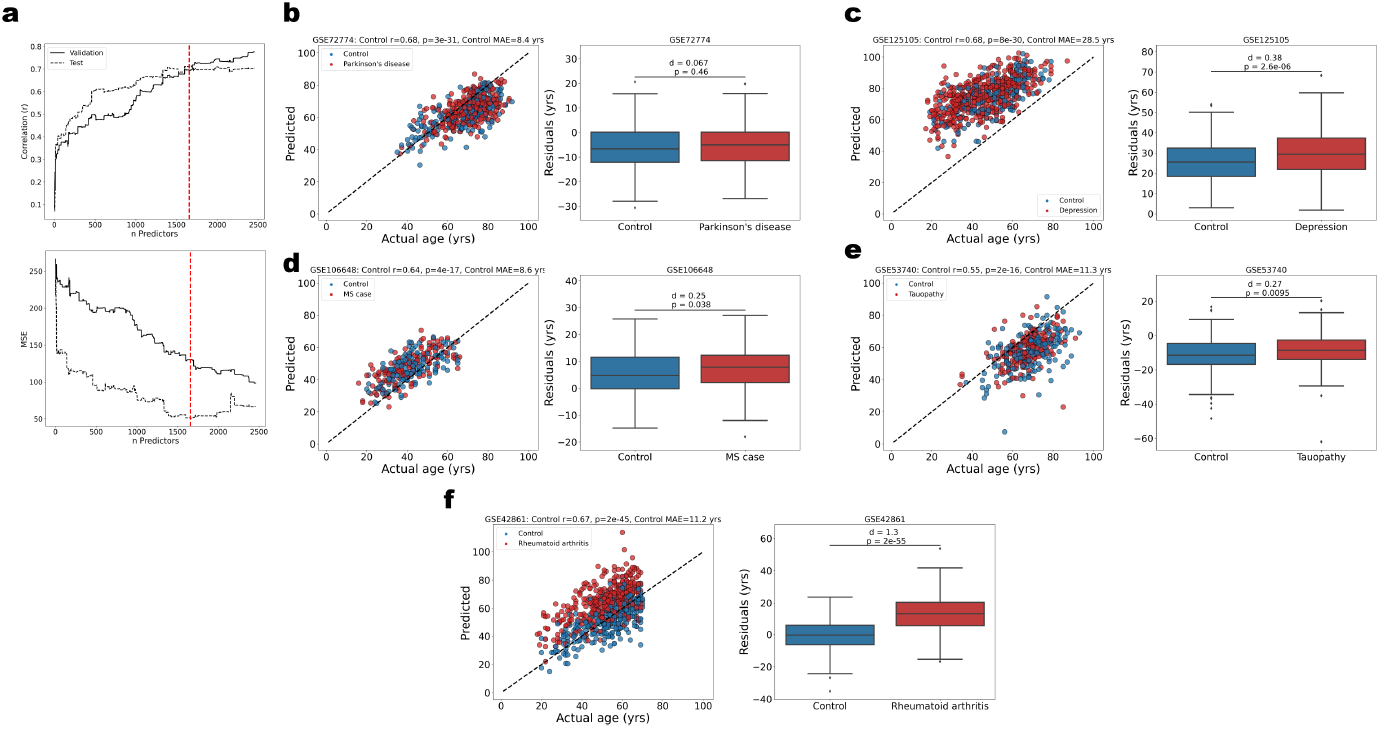
**a** (top) Correlation vs. number of features for RA-coherent mFSS model trained on features ranked by RA signal shift magnitude. (bottom) Corresponding plot for MSE vs. number of features. Red dashed line indicates number of features which give the optimum validation MSE. **b** (Left) scatter plot for predicted age vs. actual age for GSE72774. (right) box plot of the residuals of the predictions. **c** (Left) scatter plot for predicted age vs. actual age for GSE125105. (right) box plot of the residuals of the predictions. **d** (Left) scatter plot for predicted age vs. actual age for GSE106648. (right) box plot of the residuals of the predictions. **e** (Left) scatter plot for predicted age vs. actual age for GSE53740. (right) box plot of the residuals of the predictions. **f** (Left) scatter plot for predicted age vs. actual age for GSE42861. (right) box plot of the residuals of the predictions.

**Supplementary figure 6.**
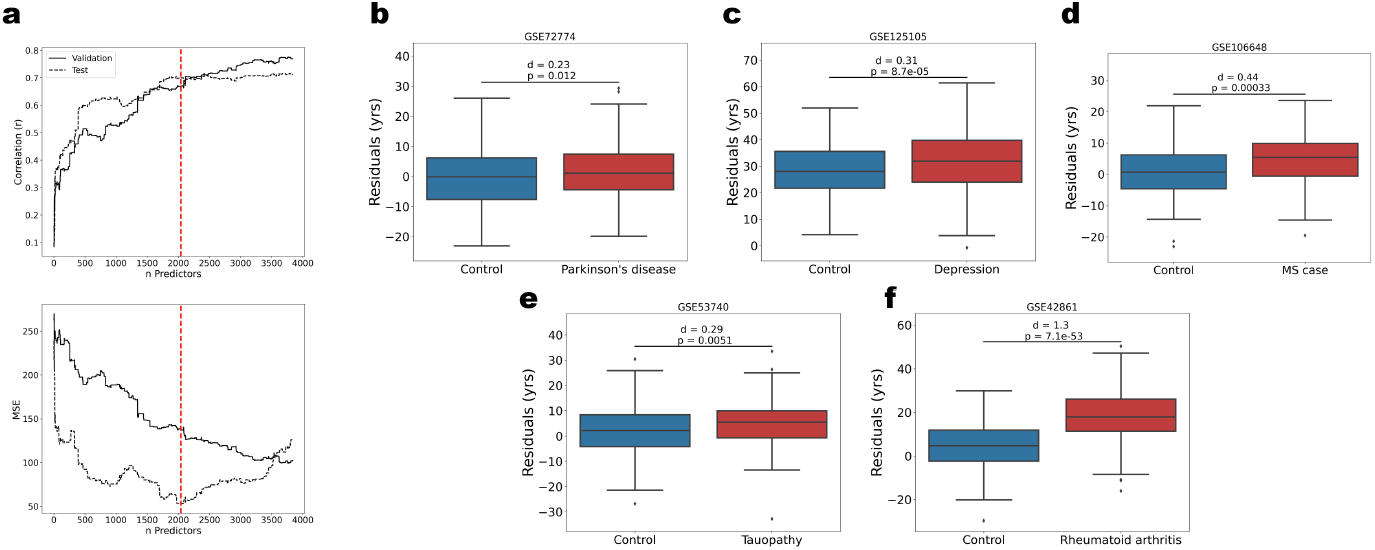
**a** (top) Correlation vs. number of features for IF-coherent mFSS model trained on features ranked by IF signal shift magnitude. (bottom) Corresponding plot for MSE vs. number of features. Red dashed line indicates number of features which give the optimum validation MSE. **b** Residuals of the IR-mFSS model predictions on GSE72774 dataset comparing healthy controls (blue) to Parkinson’s disease patients (red). **c** Residuals of the IR-mFSS model predictions on GSE125105 dataset comparing healthy controls (blue) to multiple depression patients (red). **d** Residuals of the IR-mFSS model predictions on GSE10648 dataset comparing healthy controls (blue) to multiple sclerosis patients (red). **e** Residuals of the IR-mFSS model predictions on GSE53740 dataset comparing healthy controls (blue) to tauopathy patients (red). **f** Residuals of the IR-mFSS model predictions on GSE42861 dataset comparing healthy controls (blue) to rheumatoid arthritis patients (red).

## Algorithms, Program codes and Listings

### Algorithm 1

Algorithm 1 modified Forward Stepwise Selection (mFSS) Algorithm

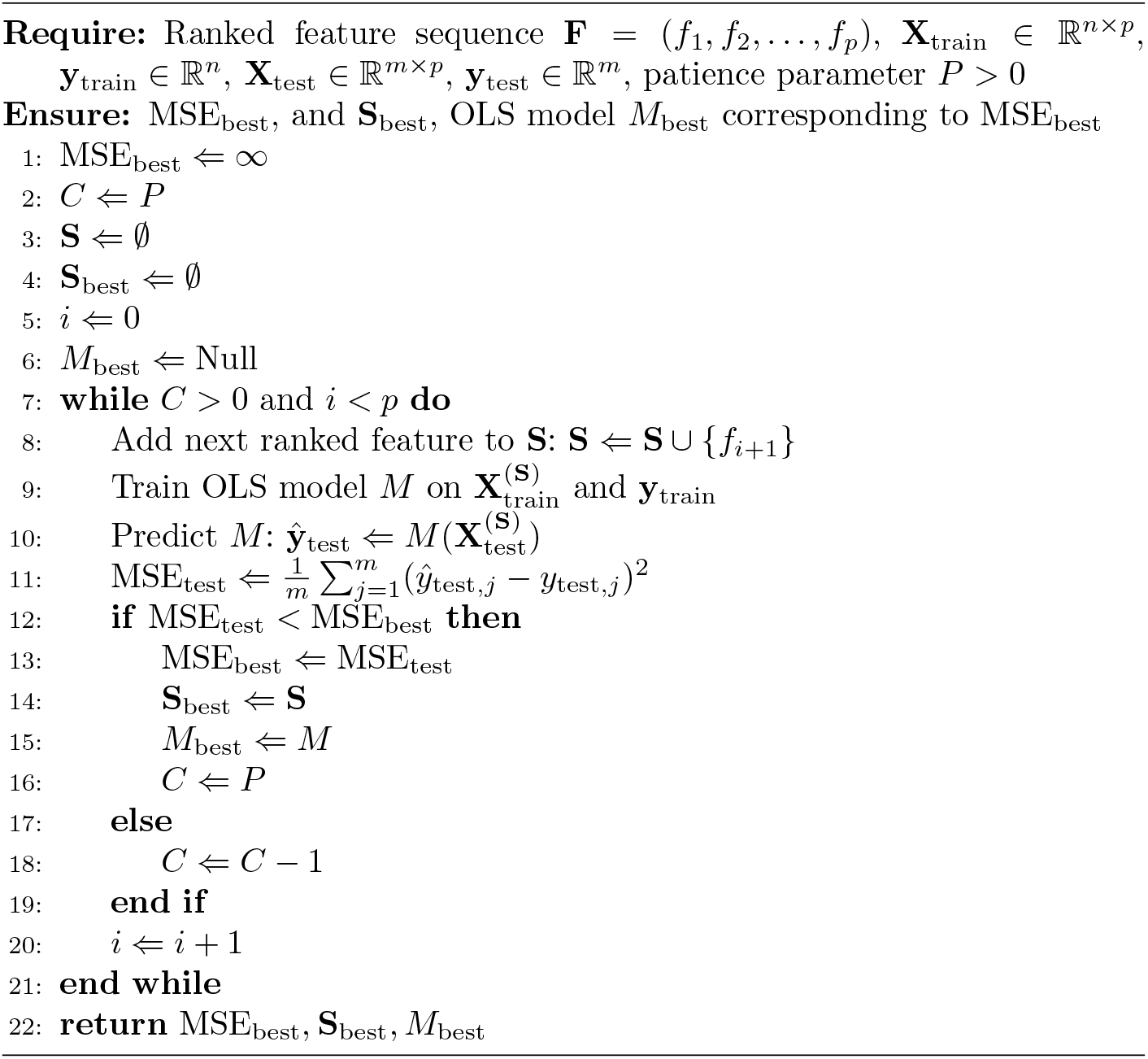

## Supplementary information

## Acknowledgments

We acknowledge and thank Dr. Anthony Hunt at UCSF for his invaluable guidance throughout the undertaking of this project, and Nathan Wong for his helpful feedback on the manuscript.

## Declarations

- Funding: NIH T32GM098218 grant and Jack Kent Cooke scholarship to CS supported this work.
- Conflict of interest/Competing interests: None
- Ethics approval: N/A
- Consent to participate: N/A
- Consent for publication: N/A
- Availability of data and materials: All data sets used in this study are publicly available on NCBI’s GEO database, found at https://www.ncbi.nlm.nih.gov/geo/.
- Code availability: The code used to process data and generate the models and figures in this study can be found at https://github.com/SkinnerCM/conboy-laboratory
- Authors’ contributions: CS wrote all code for and conducted the data analyses and model training, formulated the modified stepwise forward selection algorithm, conceived of the coherence theory for linear models and wrote the manuscript.

If any of the sections are not relevant to your manuscript, please include the heading and write ‘Not applicable’ for that section.

Editorial Policies for:

Springer journals and proceedings: https://www.springer.com/gp/editorial-policies

Nature Portfolio journals: https://www.nature.com/nature-research/editorial-policies

*Scientific Reports*: https://www.nature.com/srep/journal-policies/editorial-policies

BMC journals: https://www.biomedcentral.com/getpublished/editorial-policies

## Appendix A

## References

[1] Horvath, S.: Dna methylation age of human tissues and cell types. Genome biology 14, 115 (2013)

[2] Zou, H., Hastie, T.: Regularization and variable selection via the elastic net. Journal of the Royal Statistical Society Series B: Statistical Methodology 67(2), 301–320 (2005)

[3] Hannum, G., Guinney, J., Zhao, L., Zhang, L., Hughes, G., Sadda, S., Klotzle, B., Bibikova, M., Fan, J.-B., Gao, Y., Deconde, R.: Genome-wide methylation profiles reveal quantitative views of human aging rates. Molecular cell 49(2), 359–367 (2013)

[4] Levine, M.E., Lu, A.T., Quach, A., Chen, B.H., Assimes, T.L., Bandinelli, S., Hou, L., Baccarelli, A.A., Stewart, J.D., Li, Y., Whitsel, E.A.: An epigenetic biomarker of aging for lifespan and healthspan. Aging (albany NY) 10(4), 573–591 (2018)

[5] Lu, A.T., Quach, A., Wilson, J.G., Reiner, A.P., Aviv, A., Raj, K., Hou, L., Baccarelli, A.A., Li, Y., Stewart, J.D., Whitsel, E.A.: Dna methylation grimage strongly predicts lifespan and healthspan. Aging (albany NY) 11(2), 303–327 (2019)

[6] Belsky, D.W., Caspi, A., Corcoran, D.L., Sugden, K., Poulton, R., Arseneault, L., Baccarelli, A., Chamarti, K., Gao, X., Hannon, E., Harrington, H.: Dunedinpace, a dna methylation biomarker of the pace of aging. Elife 11, 73420 (2022)

[7] Ying, K., Liu, H., Tarkhov, A.E., Sadler, M.C., Lu, A.T., Moqri, M., Horvath, S., Kutalik, Z., Shen, X., Gladyshev, V.N.: Causality-enriched epigenetic age uncouples damage and adaptation. Nature Aging, 1–16 (2024)

[8] Horvath, S., Raj, K.: Dna methylation-based biomarkers and the epigenetic clock theory of ageing. Nature reviews genetics 19(6), 371–384 (2018)

[9] Bozack, A.K., Rifas-Shiman, S.L., Gold, D.R., Laubach, Z.M., Perng, W., Hivert, M.-F., Cardenas, A.: Dna methylation age at birth and childhood: performance of epigenetic clocks and characteristics associated with epigenetic age acceleration in the project viva cohort. Clinical Epigenetics 15(1), 62 (2023)

[10] Bell, C.G., Lowe, R., Adams, P.D., Baccarelli, A.A., Beck, S., Bell, J.T., Christensen, B.C., Gladyshev, V.N., Heijmans, B.T., Horvath, S., Ideker, T.: Dna methylation aging clocks: challenges and recommendations. Genome biology 20, 249 (2019)

[11] He, X., Liu, J., Liu, B., Shi, J.: The use of dna methylation clock in aging research. Experimental Biology and Medicine 246(4), 436–446 (2021)

[12] Bergsma, T., Rogaeva, E.: Dna methylation clocks and their predictive capacity for aging phenotypes and healthspan. Neuroscience insights 15, 2633105520942221 (2020)

[13] Stenvinkel, P., Karimi, M., Johansson, S., Axelsson, J., Suliman, M., Lindholm, B., Heimbürger, O., Barany, P., Alvestrand, A., Nordfors, L., Qureshi, A.R.: Impact of inflammation on epigenetic dna methylation–a novel risk factor for cardiovascular disease? Journal of internal medicine 261(5), 488–499 (2007)

[14] Kangaspeska, S., Stride, B., Métivier, R., Polycarpou-Schwarz, M., Ibberson, D., Carmouche, R.P., Benes, V., Gannon, F., Reid, G.: Transient cyclical methylation of promoter dna. Nature 452(7183), 112–115 (2008)

[15] Benbrahim-Tallaa, L., Waterland, R.A., Styblo, M., Achanzar, W.E., Webber, M.M., Waalkes, M.P.: Molecular events associated with arsenic-induced malignant transformation of human prostatic epithelial cells: aberrant genomic dna methylation and k-ras oncogene activation. Toxicology and applied pharmacology 206(3), 288–298 (2005)

[16] Bose, R., Onishchenko, N., Edoff, K., Janson Lang, A.M., Ceccatelli, S.: Inherited effects of low-dose exposure to methylmercury in neural stem cells. Toxicological Sciences 130(2), 383–390 (2012)

[17] Christiansen, C., Castillo-Fernandez, J.E., Domingo-Relloso, A., Zhao, W., El-Sayed Moustafa, J.S., Tsai, P.-C., Maddock, J., Haack, K., Cole, S.A., Kardia, S.L., Molokhia, M.: Novel dna methylation signatures of tobacco smoking with trans-ethnic effects. Clinical epigenetics 13, 182 (2021)

[18] Massart, F.M.S.M.P.J.E.H.J.B.-N.E.R.A.K.O.C.C.J.S.M. R., Mongrain, V.: The genome-wide landscape of dna methylation and hydroxymethylation in response to sleep deprivation impacts on synaptic plasticity genes. Translational psychiatry 4(1), 347–347 (2014)

[19] Cedernaes, O.M.E.V.S.B.J.E.V.H.D.S.L.Z.J.R.S.H.B. J., Benedict, C.: Acute sleep loss induces tissue-specific epigenetic and transcriptional alterations to circadian clock genes in men. The Journal of Clinical Endocrinology Metabolism 100(9), 1255–1261 (2015)

[20] Bhatti, Z.Y.S.X.M.K.W.S.C.L.K.K.T.H.E.A. P., Wang, P.: Nightshift work and genome-wide dna methylation. Chronobiology International 32(1), 103–112 (2015)

[21] Bind, Z.A.G.A.P.A.C.B.B.A.T.L.K.P.V.P. M.A., Schwartz, J.: Effects of temperature and relative humidity on dna methylation. Epidemiology 25(4), 561–569 (2014)

[22] McDade, R.C.P.J.M.J.H.M.K.B.J.M.G.E.K.C.W. T.W., Kobor, M.S.: Genome-wide analysis of dna methylation in relation to socioeconomic status during development and early adulthood. American journal of physical anthropology 169(1), 3–11 (2019)

[23] Mei, B.J.L.C.C.M.J. X., Conboy, I.M.: Fail-tests of dna methylation clocks, and development of a noise barometer for measuring epigenetic pressure of aging and disease. Aging (Albany NY) 15(17), 8552 (2023)

[24] Santos-Moreno, B.-A.G.M.-C.M.A.P.A.E.D.B.-N.P.K.R.A.J. P., Rojas-Villarraga, A.: Inflammaging as a link between autoimmunity and cardiovascular disease: the case of rheumatoid arthritis. RMD open 7(1), 001470 (2021)

[25] Rezus, C.A.B.A.L.A.C.C.T.B.I.S.G.D.D.N.B.C. E., Rezus, C.: The link between inflammaging and degenerative joint diseases. International journal of molecular sciences 20(3), 614 (2019)

[26] Calabrese, S.A.M.D.C.R.D.P.R.L.S.C.S.Z.M.G.J.C.E.J. V., Franceschi, C.: Aging and parkinson’s disease: Inflammaging, neuroinflammation and biological remodeling as key factors in pathogenesis. Free Radical Biology and Medicine 115, 80–91 (2018)

[27] Russo, T., Riessland, M.: Age-related midbrain inflammation and senescence in parkinson’s disease. Frontiers in Aging Neuroscience 14, 917797 (2022)

[28] Perdaens, O., Van Pesch, V.: Molecular mechanisms of immunosenescene and inflammaging: relevance to the immunopathogenesis and treatment of multiple sclerosis. Frontiers in neurology 12, 811518 (2022)

[29] Deleidi, J.M. M., Rubino, G.: Immune aging, dysmetabolism, and inflammation in neurological diseases. Frontiers in neuroscience 9, 140325 (2015)

[30] Simon, S.C.A.-H.G.P.S.H.B.C.D.W.A.S.M.V.E.M.O.J.S. M.S., Musil, R.: Monocyte mitochondrial dysfunction, inflammaging, and inflammatory pyroptosis in major depression. Progress in Neuro-Psychopharmacology and Biological Psychiatry 111, 110391 (2021)

[31] Emmert-Streib, F., Dehmer, M.: High-dimensional lasso-based computational regression models: regularization, shrinkage, and selection. Machine Learning and Knowledge Extraction 1(1), 359–383 (2019)

[32] Sawilowsky, S.S.: New effect size rules of thumb. Journal of modern applied statistical methods 8, 597–599 (2009)

